## Additional statistical analyses for "A signal of competitive dominance in mid-latitude herbaceous plant communities"

[1]Department of Applied Mathematics. Technical University of Madrid. Av. Juan de Herrera, 6. 28040 Madrid, Spain.
[2]Theoretical and Computational Ecology Group. Center for Advanced Studies (CEAB-CSIC). C. Accés Cala St. Francesc 14. 17300 Blanes, Catalonia, Spain.
[3]Universidad Autónoma de Madrid. Facultad de Ciencias Económicas y Empresariales. Depto. Análisis Económico: Economía Cuantitativa. C. Francisco Tomás y Valiente 5, 28049 Madrid, Spain
[4]Department of Bioscience. Aarhus University, Aarhus. Ny Munkegade 114, DK-8000, Aarhus C, Denmark.

February 7, 2021

### Contents

**A Intra- *vs.* interspecific strength ratio estimation for empirical data and model fitting 2**

**B AFE cell aggregation: up- and down-scaling spatial scales 2**

**C Clustering patterns across grid cells over the continent 5**

**D Robustness of the results 6**

### A Intra- *vs.* interspecific strength ratio estimation for empirical data and model fitting

The quantity $\hat{\rho}$ sets up the scale between inter- and intraspecific interactions. We cannot calculate its value directly from data, so ecoregional competition matrices were first calculated as if $\hat{\rho} = 1$. For the set of normalized height values, $t_i = (h_i - h_{\min})/(h_{\max} - h_{\min})$, we computed $\rho_{ij} = \delta_{ij} + t_j - t_i$, so that $-1 \leq \rho_{ij} \leq 1$. Maximum ($h_{\max}$) and minimum height values ($h_{\min}$) were taken over the set of realized heights in each ecoregion. Note that if $j$ is the species with largest $h$-value and $i$ is the species with smallest $h$-value and $\hat{\rho} = 1$, then $\rho_{ij} = 1$. Hence a particular interspecific interaction would be equal to the intraspecific strength ($\rho_{ii} = 1$). This is, in principle, unrealistic.

However, data from Fig. 3 permits an estimation of the ratio $\hat{\rho}$ between inter- and intraspecific interactions. Assuming $\hat{\rho} = 1$, the fitted function:

$$p_{\mathrm{c}} \sim (\langle \rho \rangle S)^{-\gamma}, \tag{A.1}$$

for empirical data is $p_{\mathrm{c}} = 7.26(\langle \rho \rangle S)^{-0.61}$. This implies that, for $\langle \rho \rangle S \sim 25.8$, $p_{\mathrm{c}} \sim 1$. A fit to model predictions in the power-law regime [see Fig. 3 for model parameters $K = 1000$, $\mu = 5$, $\alpha^{+} = 50$, $\alpha^{-} = 0.1$, and $\sigma = 0.2$, see also Capitán *et al.* (2020)] yields $p_{\mathrm{c}} = 1.02(\langle \rho \rangle S)^{-0.61}$, from which $\langle \rho \rangle S \sim 1.03$ to get $p_{\mathrm{c}} \sim 1$. To make model predictions match empirical data, the latter must be translated to the left. This amounts to a multiplicative factor $\hat{\rho}$ in the $X$ axis (note the logarithmic scale), so it is enough to choose $\hat{\rho} \approx 1.03/25.8 \approx 4 \times 10^{-2}$. Hence we obtain an indirect estimation of inter- *vs.* intraspecific overall effects. The ratio between inter- and intraspecific competition must be, on average across ecoregions, about 4%.

### B AFE cell aggregation: up- and down-scaling spatial scales

The spatial model entails signals of clustering for increasing spatial scales. We have studied the robustness of our results against different spatial resolutions using AFE plant distributions. The effect of spatial scales relies on: (i) the derivation of a species-area relationship (up-scaling), and (ii) a down-scaling to extrapolate local abundances at areas smaller than $50 \times 50$ km$^2$.

In order to derive a species-area relationship, we devised a procedure to randomly aggregate AFE cells ($50 \times 50$ km$^2$) at spatial scales larger than the original resolution of the data (Fig. B1). For a given ecoregion, we randomly chose a centroid of one of its cells and used it as a center to re-grid the ecoregion for increasing aggregated cell sizes ($100 \times 100$, $150 \times 150$ km$^2$, and so on). We then uses the new (larger) grid to define aggregated cells with increased area. For AFE cells close to the boundary of the ecoregion, the procedure simply aggregated those cells that belong to the ecoregion. For each cell aggregation we computed its area $a$ by adding the area of all the aggregated $50 \times 50$ cells, each of which was estimated as the surface of the corresponding latitude and longitude quadrangle over a sphere with radius

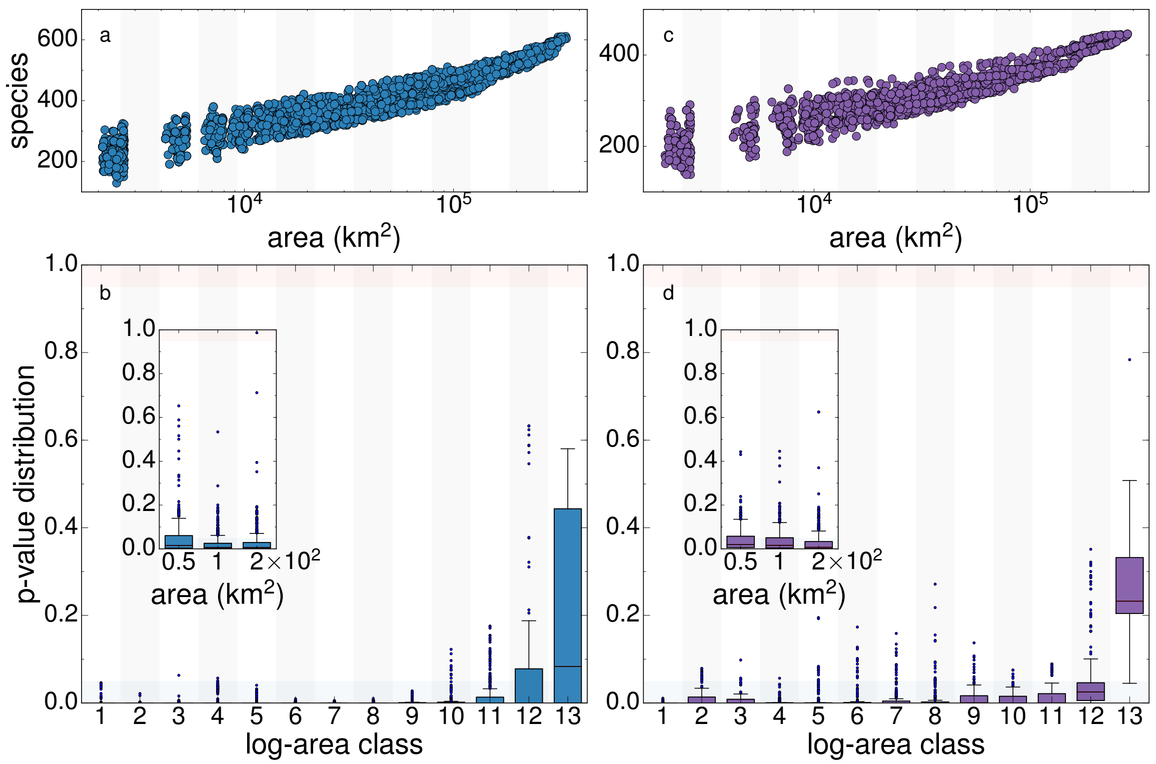

Figure B1: Clustering patterns of two ecoregions characterized by high clustering index (Atlantic mixed forests, panels **a** and **b**; Sarmatic mixed forests, panels **c** and **d**) were analyzed at increasing aggregation scales (main plots in panels **b** and **d**). As expected, as community sizes increase, the signal of significant clustering weakens. Over a critical aggregation scale, randomization tests show no signal of clustering (class 11 for Atlantic mixed forests, which corresponds to $10^5$ km$^2$; area has been divided into logarithmic bins in these panels). The insets in panels **b** and **d** show a down-scaling of the same randomization tests. Under a random placement hypothesis, communities of smaller sizes (from 50 to 200 km$^2$) were built by randomly selecting a number of species as predicted by the empirical species-area relation of the corresponding ecoregion (panels **a** and **c**). The clustering pattern robustly persists at smaller spatial scales. We reproduced Fig. 7c here (panel **b**) for completeness.

equal to the Earth's radius. The diversity $s$ of the aggregated cell is the sum of species richness reported in each $50 \times 50$ cell.

We found a clear species-area relationship (Figs. B1a and c). A linear regression of raw species richness and areas apparently did not satisfy the homoscedasticity requirement. In order for the linear regression model hypotheses to be valid, we applied a data transformation and found that for the relationship $s^\tau \sim \log a$ we could not reject the homoscedasticity condition for residuals over a range of areas (see Fig. B2). Although the hypothesis of normality was rejected by standard tests, deviations from normality are not excessively large so we used the linear regression model to extrapolate species diversities at smaller spatial scales (down-scaling). For a given area $a$, we sampled local diversities $s_i$ as $s_i = (A + B \log a + u_i)^{1/\tau}$, where $A$ and $B$ are the fitted parameters for the linear model and $u_i$ are trials of a $N(0, \sigma)$

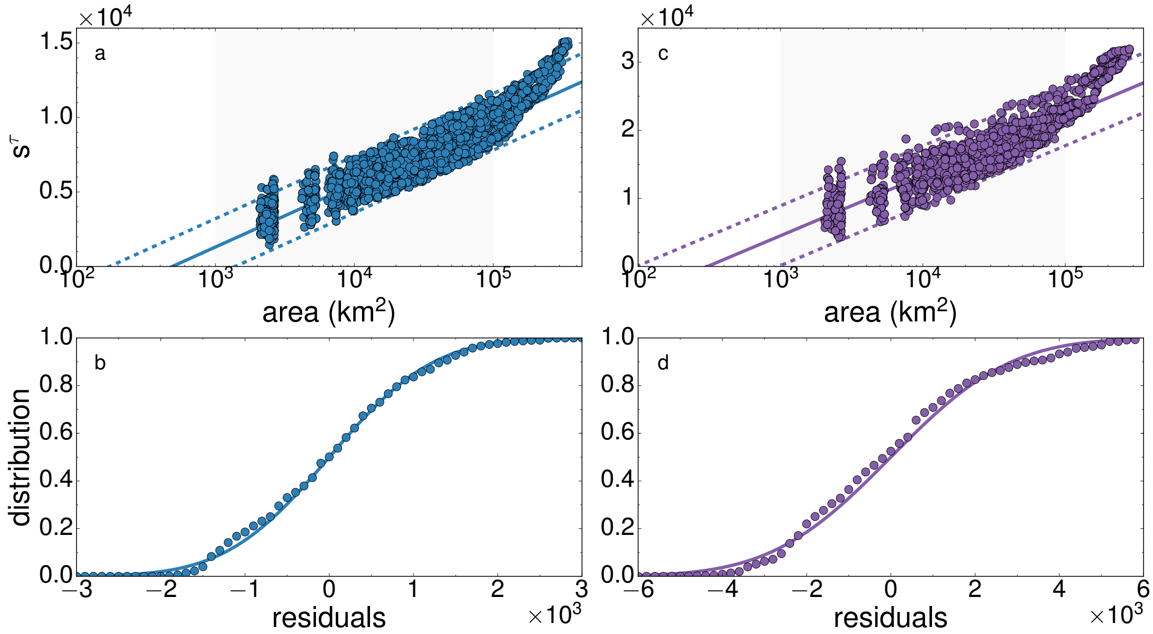

Figure B2: Diagnosis for the species-area relationship. Panels a and b refer to Atlantic mixed forests, whereas c and d summarize the diagnosis for Sarmatic mixed forests. The relationship $s^\tau \sim \log a$ is apparently linear within the area range $[10^3, 10^5]$ km$^2$ ($r^2 = 0.86$ for Atlantic and $r^2 = 0.83$ for Sarmatic mixed forests, $p < 10^{-3}$ in both cases), so the regression analysis was limited to this interval. For the first ecoregion, an exponent $\tau = 1.5$ led to a $p$-value $p = 0.36$ for the Koenker-Basset heteroscedasticity test (for Sarmatic mixed forests we got $p = 0.77$ for this test for $\tau = 1.7$), so homoscedasticity could not be rejected in both cases. 95% confidence lines are shown. Although standard tests (Kolmogorov-Smirnov) rejected the hypothesis of normality, deviations from normality are not dramatic (panels b and d) and are probably caused by a size effect (the set of areas is very large and minimal deviations from normality are detected by the test).

distribution, $\sigma$ being estimated by the standard error of regression.

Communities of smaller sizes were built up by selecting at random a $50 \times 50$ cell and randomly sampling in this cell a number $s$ of species according to the fitted species-area relationship. This procedure was repeated for a sequence of 100 values of species richness $s$; in each case $s$ species were sampled from 5 randomly $50 \times 50$ cells, which yielded 500 different realized communities at a given spatial scale. Under this random replacement hypothesis, we recovered the clustering pattern at smaller scales (Fig. B1, insets). Since individual plant interactions occur at small spatial scales, the role of species interactions in driving any community pattern should potentially increase as the scale definition of the community shrinks, so the pattern should become more conspicuous the lower the spatial scale is. Since our randomization tests compare realized diversity in grid cells to *random samples* taken from the set of species reported in the whole ecoregion, if clustering is reported at a large scale, then on average we expect clustering as well at smaller scales, even if species are chosen at random to define communities at smaller sizes. Our expectation that clustering should be more conspicuous at smaller scales is also a consequence of the spatially-explicit model [see Fig. 4 in Capitán *et al.* (2020), where model communities keep clustered in heights when the grid size decreases]. Indeed, let us consider, for example, a grid size $\ell < L$, $L$ being the size of the whole lattice (in the data, $\ell$ will correspond to the 50 km distance). If height clustering is reported at size $\ell$ is because height differences are significantly smaller than average height differences in the whole lattice. When down-scaling to sizes smaller than $\ell$ we are choosing among a subset of species which are actually clustered (recall that we always compare to the potential diversity in randomization tests), so we must recover clustering patterns even if smaller communities are defined by randomly sampling the species present in the cell of size $\ell$.

In addition, we scaled our communities down under a random placement hypothesis and recovered the same clustering pattern, so the strength of the pattern should work as a lower bound to what we may expect in reality (i.e., clustering patterns should be more intense in non-random, realized communities at smaller spatial scales).

### C Clustering patterns across grid cells over the continent

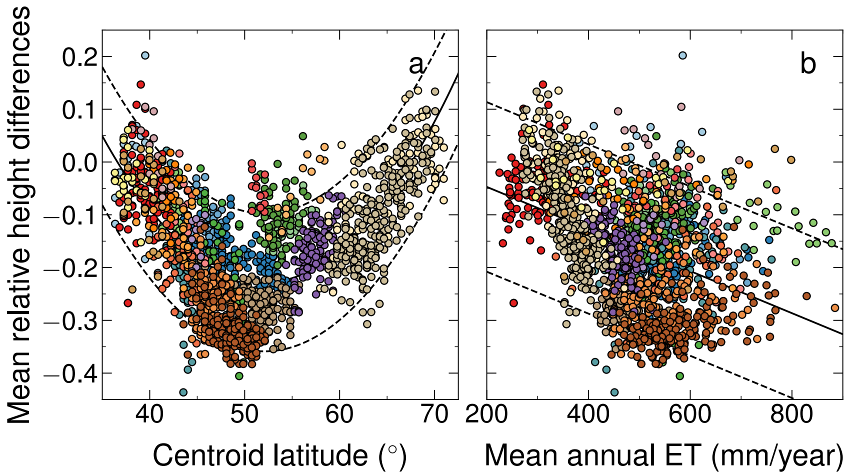

Figure C1: Clustering patterns in maximum stem height (as measured by mean height differences relative to ecoregion averages) are represented against the centroid latitude and the mean annual ET associated to each grid cell (panel **a**, quadratic fit: $r^2 = 0.45$, $p < 10^{-3}$: and **b**, linear regression: $r^2 = 0.17$, $p < 10^{-3}$, respectively; 95% confidence lines are shown). Colors used for data match codes in Fig. 2 form the main text. Species height differences show also a $U$ pattern in terms of latitude (compared to the inverse $U$ pattern in Fig. 5a), and decrease as actual evapotranspiration increases (compare to the increasing trend in Fig. 5c). At the cell level, the intensity of clustering in species maximum stem height increases as evapotranspiration increases even within the same ecoregion (better seen for those spanning higher latitudinal ranges).

### D Robustness of the results

In this Section we evaluate the robustness of our results when some assumptions are relaxed. We here show that the main conclusions remain unchanged when: (i) heights are not log-transformed; (ii) species selection in randomization tests is weighted according to species dispersal abilities; and (iii) variability in height data is allowed.

#### D.1 Log-transformation

First, our results regarding height clustering and its relation with latitude and mean annual evapotranspiration are consistently recovered when clustering indices are calculated using raw height values (without applying the logarithmic transformation) for herbaceous plants. The analysis of robustness and its results are summarized in Fig. D2. The amount of explained variance for each fit (detailed in the caption) should be compared to the values reported in the main text.

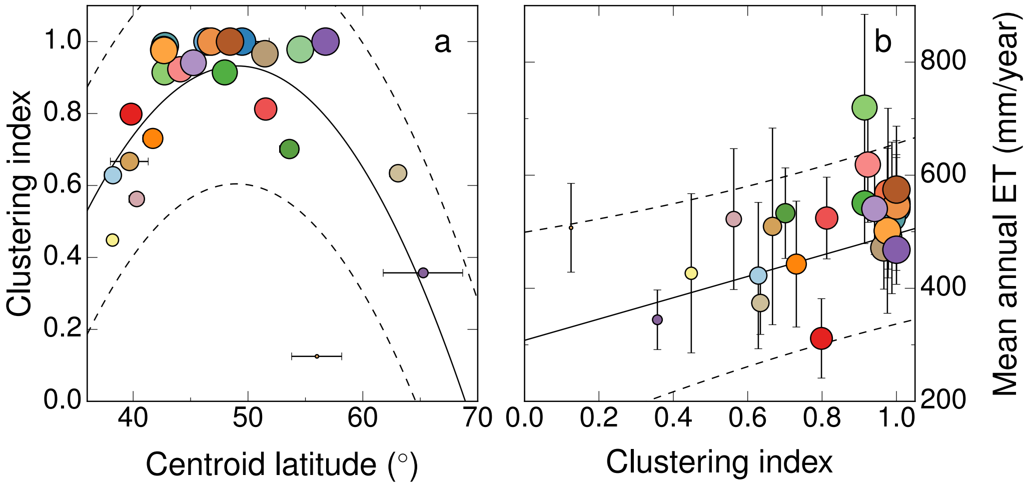

Figure D2: This figure is the counterpart of Fig. 5 (panels a and c, main text) when the logarithmic transformation is not applied in the calculation of competition values. (a) An unweighted fit to a quadratic function (least squares polynomial fit) yields $r^2 = 0.40$ with $p = 0.011$ for the dependence with latitude. (b) A weighted linear regression, with $r^2 = 0.27$ and $p = 0.03$, is here obtained for the relation between mean annual evapotranspiration and clustering index. The radius of each circle is proportional to its clustering index, and the 95% prediction curves are depicted.

#### D.2 Dispersal

Our randomization tests draw random assemblages of species from the ecoregion. Other null models for community assembly have been proposed, though (Chase *et al.*, 2011). Species can be sampled from distributions other than uniform. To test the robustness of the results based on uniform species sampling, we have conducted randomization tests that incorporate an ecologically meaningful property in the selection of sampling weights.

One of the underlying assumptions we made when partitioning ecoregions into communities is that no significant dispersal occurs between cells, and the only source for species in those communities is the ecoregion pool. Instead of uniformly sampling the ecoregion, it is more realistic to draw species in terms of their ability to colonize or, equivalently, in terms of their dispersal abilities. In the absence of reliable data for species dispersal rates, we used average seed masses as a proxy for species dispersal abilities. Data for average seed masses $w_i$ were collected from the same sources than maximum stem height. In order to sample species with high dispersal abilities, which will be better colonizers, we selected them according to two probability densities, one for which the weight assigned to species $i$ was proportional to $e^{-w_i}$, and the second density where the probability was proportional to $w_i^{-1}$. In both cases, the larger the seed mass the lower the dispersal ability of the species.

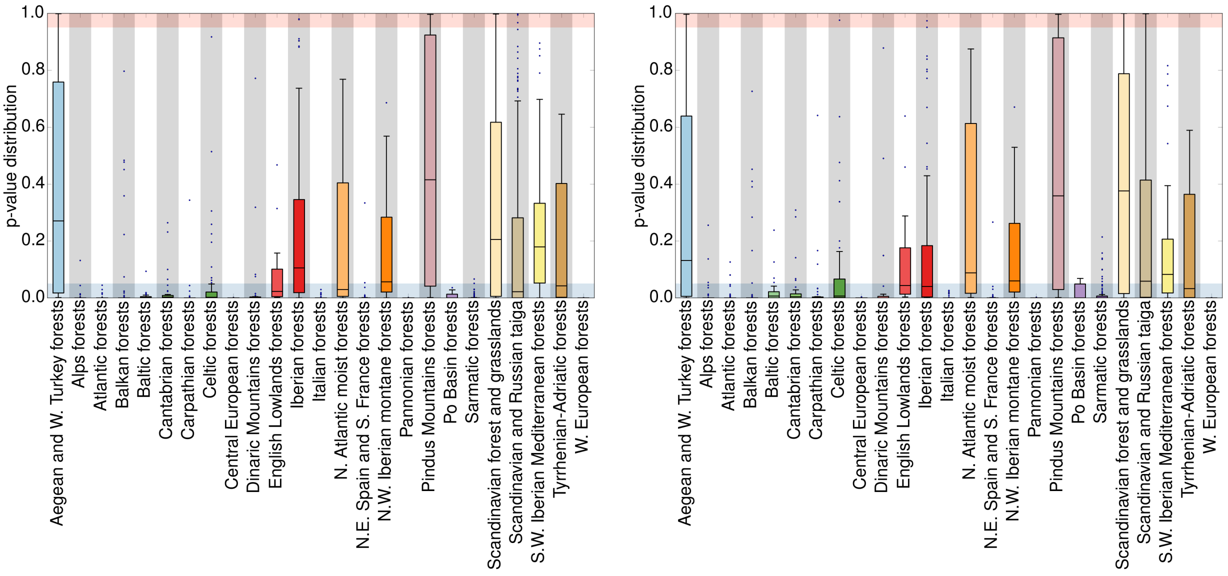

Figure D3: Left: $p$-value distributions yielded by randomization tests weighted by species seed masses $w_i$ as $e^{-w_i}$. Results are very similar to those obtained by uniform, dispersal-independent sampling (Fig. 4 of the main text). Right: weights are taken as $w_i^{-1}$. Distributions are clearly displaced to the region of significant clustering. Refer to Fig. 4 of the main text for the color code used.

Non-uniform, dispersal-weighted randomization tests yield results similar to that of unweighted tests (compare the boxplots depicted in Fig. D3 with those of Fig. 4 of the main text). The null model for community assembly based on exponential weights produces almost indistinguishable results, whereas the $w_i^{-1}$ weighting scheme results in high levels of clustering in most of mid-latitude ecoregions, although some of them depart (not dramatically) from the trend predicted by a uniform weighting procedure. In the latter case, the distributions that lied originally within the significant clustering band but now exhibit a wider boxplot, are asymmetrically biased to the region of significant clustering (Fig. D3, right panel). In summary, extended randomization tests, which sample communities according to species dispersal abilities, produce predictions comparable to that of uniform sampling. This reinforces the robustness of our conclusions.

#### D.3 Variability in heights

Height is an enormously plastic trait. The fact that different species in the trait database are likely to have been measured under different conditions may add substantial observation error to our analyses. Potential measurement errors can eventually propagate and lead to additional errors in our clustering patterns. We cannot get rid or these potential biases unless changing to improved trait datasets, but we can however test the robustness of our results using synthetic height data. In particular, we have designed a new numerical experiment in which variability (above or below) is allowed in plant heights, and repeated our analyses to check whether the clustering patterns remained or blurred. Fortunately, our results are very robust to large changes in plant height. We recalculated panels a and c of Fig. 5 (main text) but admitting up to a 20% variability in plant heights, see Fig. D4. The latitudinal pattern of clustering index, as well as the correlation with mean actual evapotranspiration, are very robust to this level of variability.

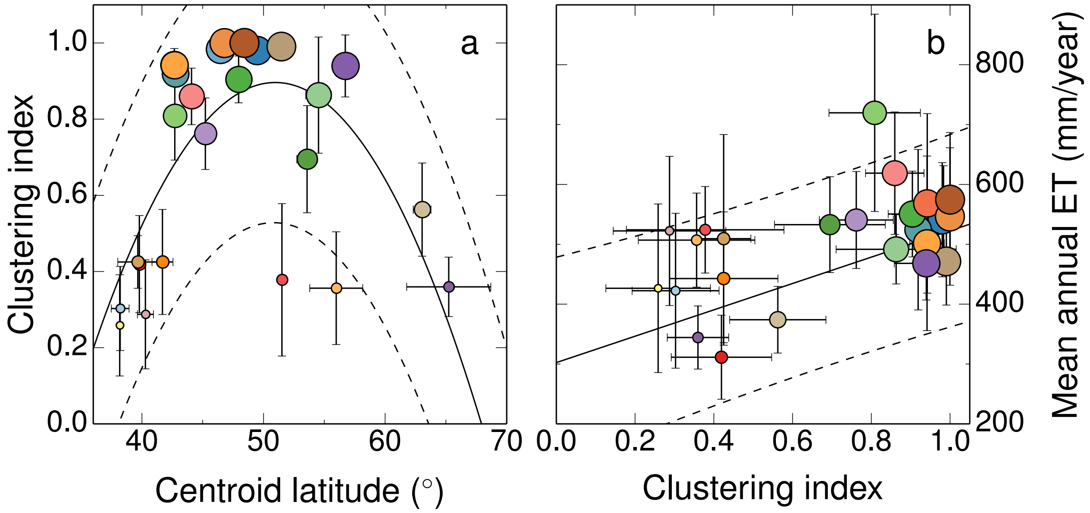

Figure D4: Counterparts of panels a and c of Fig. 5 (main text) obtained by averaging over 50 realizations of synthetic sets of heights allowing up to a 20% variability. Goodness of fit: $r^2 = 0.46$, $p = 0.004$ for the dependence with latitude (quadratic fit, panel a); $r^2 = 0.34$, $p = 0.011$ for the relation clustering index vs. ET (linear regression, panel b). Refer to the main text for the color code used.

This figure was obtained by averaging over 50 realizations of the set of heights, by admitting variability in height values to generate a synthetic set of heights in each realization. For that purpose, we draw the variables $z_i \sim N(0,1)$ and defined new heights as $\hat{h}_i = (1 + 0.2z_i)h_i$ (provided that the combination $1 + 0.2z_i$ is always positive), thus allowing up to a 20% variability in synthetic heights $\hat{h}_i$. Clustering indices change over realizations, which leads to a vertical error bar in panel a (horizontal in panel b) not present in Fig. 5 (main text). Compare with panels a and c of Fig. 5. The effect of plant height variability affects mildly to ecoregion clustering, which is an averaged measure across all locations and species reported in the ecoregion. Therefore, in spite of the large maximum level of admitted variability, we are confident that potential errors in reported plant height do not mirror into our main result, which remains almost unchanged: observed communities remain significantly clustered in plant height in mid-latitude ecoregions. This leads us to think that probably the particular set of species that are present in each cell are key to the observational pattern, instead of specific and precise values of plant height.
